# IL-6 is dispensable for causing cachexia in the colon carcinoma 26 model

**DOI:** 10.1101/2023.05.02.539076

**Authors:** Young-Yon Kwon, Sheng Hui

## Abstract

Various cytokines have been implicated in cancer cachexia. One such cytokine is IL-6, which has been deemed a key cachectic factor in mice inoculated with the colon carcinoma 26 (C26) cells, one of the most widely used models of cancer cachexia. Here to test the causal role of IL-6 in cancer cachexia, we used CRISPR/Cas9 editing to knock out *IL-6* in C26 cells. We found that growth of IL-6 KO C26 tumors was dramatically delayed. Most strikingly, while IL-6 KO tumors eventually reached the similar size as wild-type tumors, cachexia still took place, despite no elevation in circulating IL-6. We further showed an increase of immune cell populations in IL-6 KO tumors and the defective IL-6 KO tumor growth was rescued in immunodeficient mice. Thus, our results invalidated IL-6 as a necessary factor for causing cachexia in the C26 model and revealed instead its important role in regulating tumor growth via immune suppression.

## Introduction

Cancer cachexia is a systemic wasting syndrome prevalent in cancer patients, contributing to physical disability, poor quality of life, and death^1–4^. In search for mediators of cancer cachexia, researchers have proposed many cachectic factors over the years, with more than 20 of them listed in a recent review^5^. Targeting these factors, however, has so far not yielded an effective treatment. While this lack of success in translation may arise from differences between preclinical models and humans, it is possible that the real biological role of these factors has not been sufficiently established in preclinical models.

More than half of the proposed cachectic factors are cytokines^5^, with well-known examples as TNF-α^6–8^, IL-1^9,10^, IFN-γ^11,12^ and IL-6^13,14^. Among these, IL-6 is one of the most cited cachectic factors, first proposed in the syngeneic tumor model of colon 26 carcinoma (C26)^13^ and later implicated in various cancer models including genetically engineered mouse models such as the *Apc^Min/+^* colorectal cancer model and the pancreatic ductal adenocarcinoma (PDAC) model^15,16^ and patient-derived tumor xenograft models such as the HT1080 fibrosarcoma model, the TOV21G ovarian cancer model, and the A2058 melanoma model^17–19^. Given the prominent status of IL-6 in cancer cachexia research, here in our study we aim to closely examine its causal role in cancer cachexia. For this, we focus on the C26 model, where IL-6 was first proposed to be a cachectic factor^13,14^ and its cachectic role seems to be best established.

Existing evidence shows that IL-6 is unlikely a sufficient factor for causing cachexia in the C26 model. Although circulating levels of IL-6 were dramatically increased in mice inoculated with cachectic C26 cells, similar level of IL-6 was also measured in mice inoculated with a non-cachectic subclone of C26 cells^20,21^, indicating that IL-6 alone cannot cause cachexia. Furthermore, infusion of IL-6 to healthy and non-cachectic C26 tumor-bearing (TB) mice failed to induce body weight loss^22,23^. Although the infusion of IL-6 to mice injected with CHO cells led to weight loss and muscle wasting, the circulating IL-6 levels (80-100ng/ml) were several orders of magnitude higher than the typical level (0.1-1ng/ml) measured in cancer cachexia models^24^, which prevents it as strong evidence for supporting IL-6’s sufficiency in cachexia.

Unlike sufficiency, the necessity of IL-6 in causing cachexia in the C26 model seems supported by clear evidence. First, studies have shown neutralizing circulating IL-6 with anti-IL-6 antibody partially prevented the body weight loss^13,23,25^. Second, as the cancer cells are the main source of circulating IL-6 in the C26 model^26^, the inhibition of IL-6 expression in the C26 cells by shRNA dramatically lowered the circulating IL-6 level and almost completely prevented body weight loss^27^. Based on these results, IL-6 was concluded to be necessary for causing cachexia in the C26 model. However, surprisingly, there has been no study that completely disrupts the IL-6 expression by the C26 tumor and determines its effect on tumor and host. Unlike the antibody treatment and the shRNA knockdown experiments described above, such an experiment would create a condition where IL-6 remains at basal level, thus serving as a more rigorous test on whether IL-6 is necessary for causing cachexia.

In this study, we tested the necessity of IL-6 in cachexia in the C26 model by knocking out the *IL-6* gene in the C26 cells using CRISPR and closely monitoring tumor growth and body weight. We found that IL-6 KO significantly slowed tumor growth, and while the IL-6 KO tumor eventually grew in size cachexia still took place, creating a situation where cachexia happens without elevated circulating IL-6, directly against IL-6 being necessary for causing cachexia in the C26 model. We further demonstrated that IL-6 promotes tumor growth via repressing immune infiltration in tumors. Moreover, with the same approach, we found that the other proposed cachectic factor in the C26 model, leukemia inhibitory factor (LIF), is not necessary for causing cachexia in the C26 model either.

## Results

### Confirmation of the correlation between elevated circulating IL-6 and cachectic phenotypes in the C26 model

To reveal the real role of IL-6 in cachexia in the C26 model, we first aimed to test the observation that circulating IL-6 level is strongly correlated with cachectic phenotypes in the C26 model^13,15,19^. For this, in addition to a well-established cachectic C26 cell line (cxC26)^28^, we used a non-cachectic C26 cell line (ncxC26) as a control. In the model, cells are subcutaneously injected into syngeneic host CD2F1 mice. To control for differences in growth rates between the two cell lines, half a million ncxC26 cells were injected while a million cxC26 cells were used due to faster growth of ncxC26 than cxC26 in the host. Within 2 weeks post tumor implantation, the mice bearing the cxC26 tumor lost more than 15% body weight loss, while the mice bearing the ncxC26 tumor maintained body weight (Figure 1a), despite similar tumor progression (Figure 1b). Consistent with body weight loss, the lean and fat mass were decreased only in the cxC26 group (Figure 1c and 1d). This is reflected in smaller white and brown adipose tissues, and quadriceps and heart (Figure 1e), as well as declined muscle function (Figure 1f) in the cxC26 group. Interestingly, the kidney size was slightly reduced and, the spleen size doubled in cxC26 TB mice, whereas liver mass was not altered (Figure 1e).

**Fig. 1.**
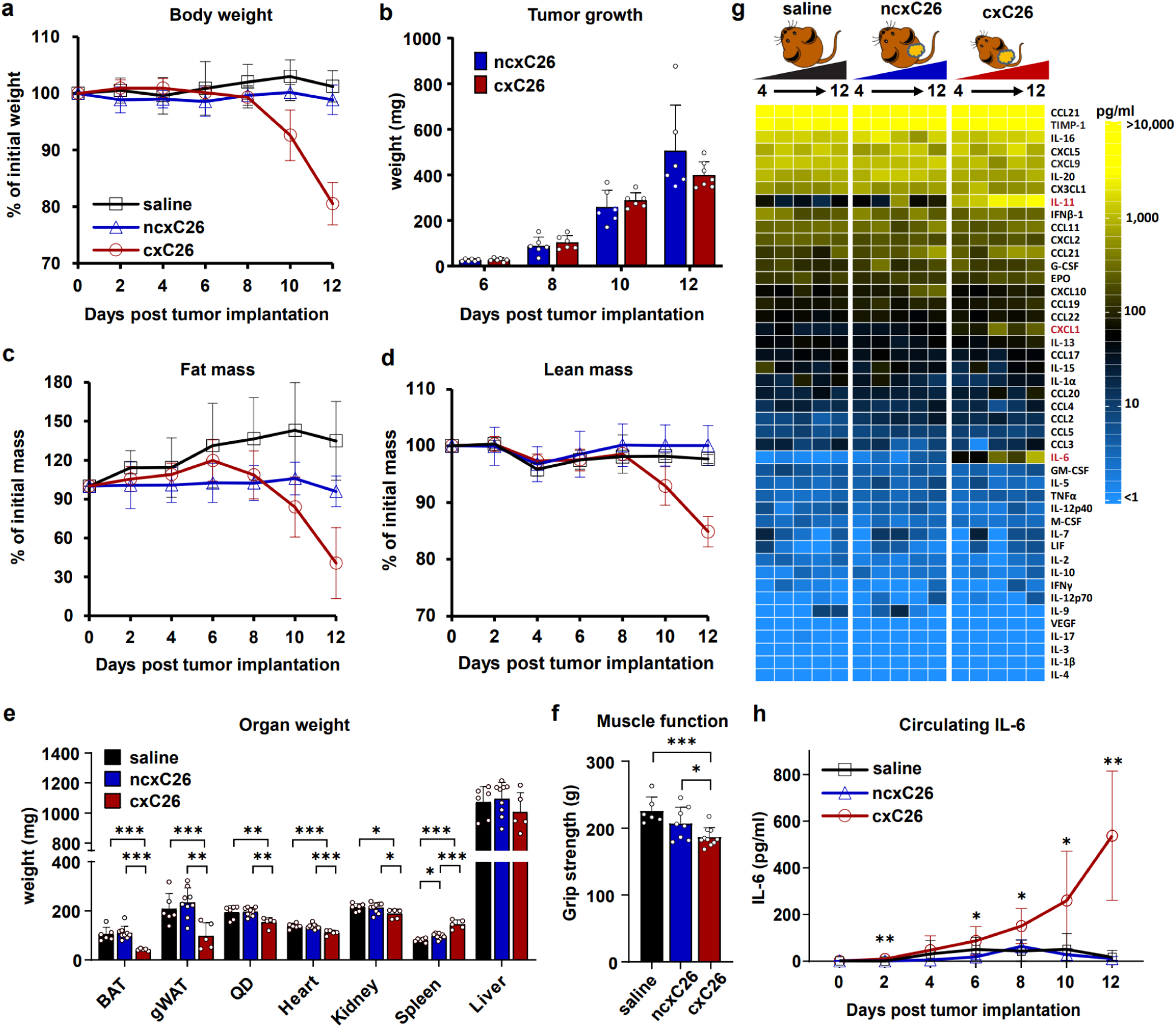
Elevation of circulating IL-6 in mice bearing the cachectic C26 (cxC26) tumor but not in those bearing the non-cachectic C26 (ncxC26) tumor. CD2F1 male mice were inoculated with 1 × 10^6^ cxC26 or 0.5 × 10^6^ ncxC26 cells. **a** Body weight, **b** tumor mass, **c** fat mass, and **d** lean mass post-implantation for mice injected with saline, cxC26 cells, or ncxC26 cells. n = 6 for saline, n = 8 for ncxC26, n = 12 for cxC26. The fat and lean mass were measured using Echo-MRI. **e** Mass of brown adipose tissue (BAT), gonadal white adipose tissue (gWAT), quadriceps (QD), heart, kidney, spleen, and liver on day 12 post implantation. n = 6 for saline, n = 9 for ncxC26, n = 5 for cxC26. **f** Muscle function of mice as evaluated using a grip strength meter. n = 6 for saline, n = 7 for ncxC26, n = 8 for cxC26. **g** Circulating cytokine profiles in saline, cxC26, and ncxC26-bearing mice. Mice were euthanized every two days from day 4 to day 12 for blood collection. The cytokine profiles were measured using Luminex 44 cytokine panel. n = 5∼6 biologically independent animals per group and time point. Cytokines highlighted in red are those with significantly altered levels in cxC26 compared to ncxC26 and saline (adjusted p-value < 0.01 by two-way ANOVA). **h** Circulating IL-6 concentration measured using a Proquantum immunoassay. n = 7 per group. All data in the figure are shown as the mean ± s.d. Significance of the differences: *\*P < 0.05, ** P < 0.01, *** P < 0.001* between groups by one-way ANOVA (**e, f**) or two-way ANOVA (**b, h**).

With the model and its control established in our hands, we went on to measure the circulating levels of IL-6 and other cytokines using a 44-cytokine Luminex panel. The measurement was done for 6 plasma samples collected every other day along the disease progression. The result showed that the circulating level of IL-6 indeed increased over time in the cxC26 group, with significantly higher level already in day 4 and day 6 even before the onset of body weight loss (Figure 1g). Interestingly, among the only other two significantly elevated cytokines in the panel is IL-11 (Figure 1g), another member of the IL-6 cytokine family. Note that both IL-6 and IL-11 were excessively released by cxC26 cells under a stress condition compared to ncxC26 cells (Figure S1), suggesting tumor as an important source for them. To further confirm the elevation of IL-6 in the cxC26 group, we measured circulating IL-6 levels using a highly sensitive immunoassay, and found that circulating IL-6 was even elevated within 2 days post tumor implantation (Figure 1h). Together, these results confirmed the strong correlation between circulating IL-6 level and cachexia in the C26 model. Next, we wished to test the causality of circulating IL-6 in cancer cachexia.

### Knockout of *IL-6* in the C26 cells delays the progress of cachexia only because it inhibits tumor growth

To test whether blocking the secretion of IL-6 by cachectic tumor prevents body weight loss, we set out to construct *IL-6* knockout using CRISPR/cas9 in the cxC26 cells. We first obtained the cxC26 IL-6 knockout pool (IL-6 KOp), which reduced 86% and 95% of the secretion of IL-6 under normal and stress conditions in cell culture, respectively (Figure S2a). Inoculated with the IL-6 KOp cells, mice showed a ∼2-week delay in body weight loss, consistent with a previous study^27^ (Figure S2b). Surprisingly however, the growth of the IL-6 KOp tumor was substantially slower than that of the wild-type cxC26 tumor (Figure S2c). In fact, the IL-6 KOp tumor was barely detectable when the wild-type cxC26 tumor already caused body weight loss (Figure S2c). As this growth defect was not observed in vitro (Figure S2d), the results indicate IL-6’s important role in tumor growth in vivo, casting doubt on its role in cachexia.

Next, to further test the role of IL-6 in cachexia, we completely disrupted the IL-6 secretion by the cxC26 tumor by isolating the two subclones (s1 and s2) of cxC26 IL-6 KO cells (Figure 2a). Consistent with the IL-6 KOp, IL-6 KO s1 showed severe growth defect. With the injection of 1 million IL-6 KO s1 cells, the tumor grew slowly in the first week, and then shrunk and disappeared one month post injection. We then injected 10 million cxC26 IL-6 KO s1 cells and with a slow initiation period the tumor started to grow after 3 weeks (Figure 2b). With a similar tumor size at the end point, the cxC26 IL-6 KO s1 tumor caused similar body weight loss and muscle wasting as the wild-type cxC26 tumor (Figure 2b, c, d and e). This is striking because at the end point when cachexia took place, there was no elevation of circulating IL-6 in the cxC26 IL-6 KO s1 group compared to non-tumor group (Figure 2f), pointing to factors other than IL-6 as mediators of cachexia here. These observations were also observed in the cxC26 IL-6 KO s2 cell line (Figure S3).

**Fig. 2.**
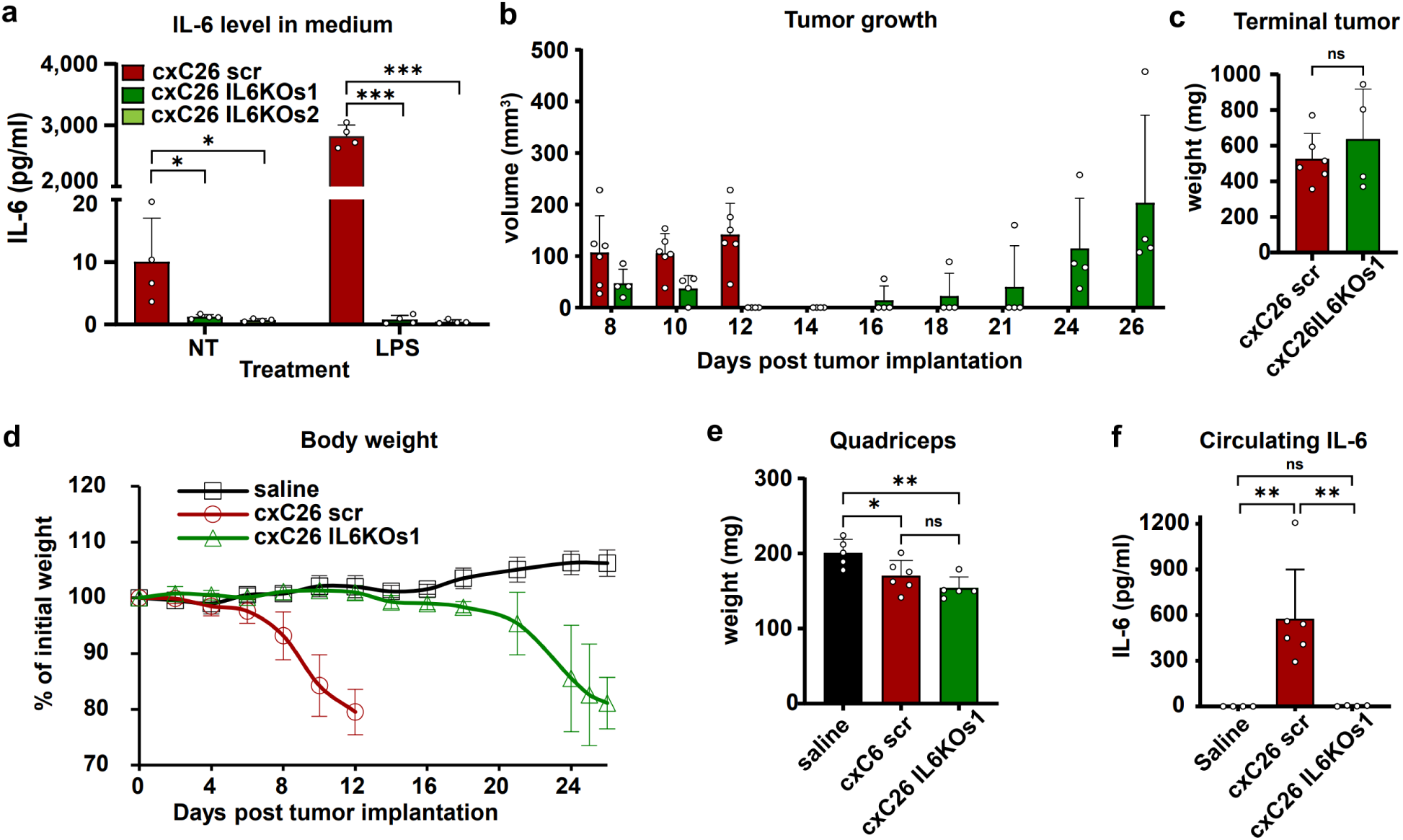
Disruption of the *IL-6* gene in cachectic C26 cells delays tumor growth, but still leads to body weight loss without elevation of circulating IL-6. **a** IL-6 level in conditioned media of cxC26 scr (CRISPR/Cas9 scrambled gRNA control) and cxC26 IL-6 KO subclones 1 (s1) and 2 (s2) with or without treatment of lipopolysaccharide (LPS) for 24 hours. (n = 4 per group) **b-f** CD2F1 male mice were inoculated with 1 × 10^6^ cxC26 scr or 1 × 10^7^ cxC26 IL-6 KO s1 cells. **b** Tumor growth and **c** body weight in saline, cxC26 scr or cxC26 IL-6 KO s1 tumor-bearing mice. **d-f** All groups were euthanized when the tumor-bearing mice developed cachexia (more than 15% of body weight loss). **d** Tumor weight, **e** muscle mass, and **f** circulating IL-6 concentration at terminal time point in saline, cxC26 scr and cxC26 IL-6 KO s1 tumor-bearing mice. **b,c,d,f** n = 4 for saline and for cxC26 IL-6 KO s1, n = 6 for cxC26. **e** n= 5 for saline and cxC26 IL-6 KO s1, n = 6 for cxC26. All data in the figure are shown as the mean ± s.d. Significance of the differences: *\*P < 0.05, ** P < 0.01, *** P < 0.001* between groups by one-way ANOVA.

To test whether these findings are applicable in the Balb/c mouse from which the original C26 cells were derived, we implanted Balb/c mice with 1 and 10 million of cxC26 and cxC26 IL-6 KO s1, respectively. Consistent with our results in the CD2F1 mice, the cxC26 IL-6 KO s1 showed slower tumor growth and bodyweight loss without elevation of circulating IL-6 (Figure S4). Altogether, these results demonstrate that IL-6 is not necessary for causing cachexia in the C26 model. The observed prevention of body weight loss upon IL-6 inhibition is simply due to impaired tumor growth in the absence of IL-6.

### Knockout of *LIF* or *IL-11*, another two IL-6 family cytokines, also inhibits tumor growth

Recent studies pointed to leukemia inhibitor factor (LIF), as another cachectic factor in the C26 model, based on the key observation that the disruption of *LIF* in tumor prevented the body weight loss^29,30^. Although we did not measure an elevation of LIF by the cytokine Luminex panel in our C26 model perhaps due to the very low concentration of circulating LIF (Figure 1g), we observed a significantly higher secretion of LIF in cxC26 cells compared to ncxC26 cells (Figure S1). To test the causality of LIF in cachexia in the C26 model, we monitored body weight and tumor growth using another cachectic C26 cell line (referred to as C26nci as it was obtained from the National Cancer Institute) and its LIF KO subclone. The C26nci and its LIF KO were used in the study that showed the cachectic effect of LIF^29^. Consistent with that study, the body weight of the C26nci LIF KO group was not significantly dropped, whereas the C26nci group showed a ∼20% of body weight loss on week 3. However, while monitored for two more weeks, mice bearing the C26nci LIF KO tumor showed clear body weight loss and muscle wasting, to the same extent as the C26nci group at the end point (Figure 3a and b). Tumor growth data showed that the disruption of *LIF* inhibited the tumor growth (Figure 3c), similar to the effect of the IL-6 disruption. Importantly, the level of circulating LIF or IL-6 was not elevated in cachectic mice bearing the C26nci LIF KO tumors (Figure 3d). To test whether these results hold for the cxC26 cell line, we constructed a CRISPR pool KO of *LIF* in the cxC26 cells, and also observed delayed body weight loss and slowed tumor growth in mice injected with the cxC26 LIF KOp cells (Figure S5). Thus, similar to IL-6, LIF mediates cachexia via its effect on tumor growth, and is not necessary for causing cachexia in the C26 model. Moreover, the observation of cachexia without IL-6 elevation using another C26 cell line provides further support to our conclusion that IL-6 is not necessary for causing cachexia in the C26 model.

**Fig. 3.**
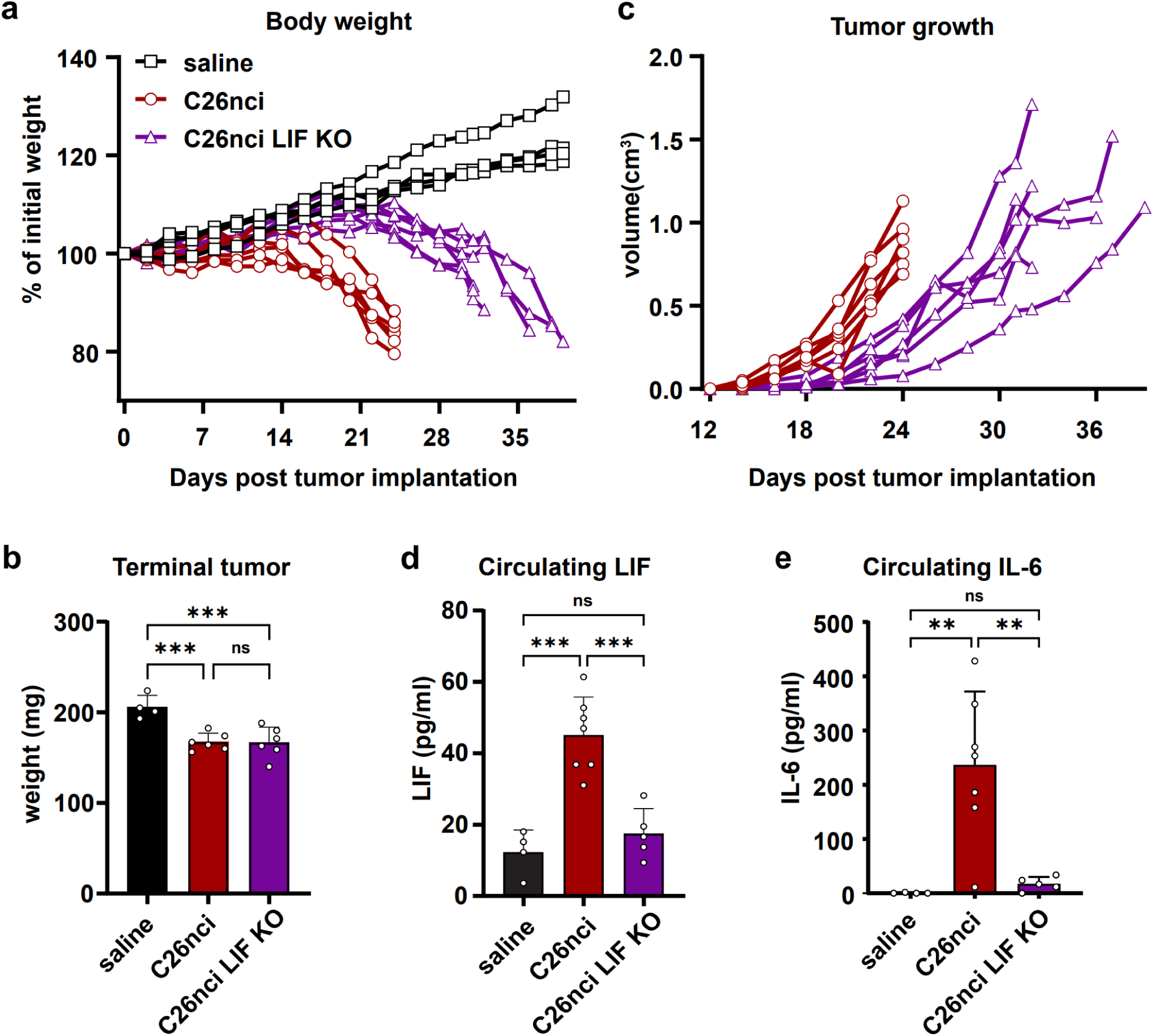
Disruption of the *LIF* gene in cachectic C26 cells delays tumor growth, but still leads to body weight loss without elevation of circulating LIF or IL-6. CD2F1 Mice were inoculated 1 × 10^6^ C26nci or C26nci LIF KO cells. All groups were euthanized when they developed cachexia phenotype (>15% of body weight loss). **a** Body weight and **b** mass of quadriceps at terminal time point. **c** Tumor growth. **a-c** n = 5 for saline, n = 6 for C26nci and C26nci LIF KO **d** Concentration of circulating LIF. **e** Concentration of circulating IL-6. **d, e** n = 4 for saline, n = 7 for C26nci, n = 5 for C26nci LIF KO. All data in the figure are shown as the mean ± s.d. Significance of the differences: *\*P < 0.05, ** P < 0.01, *** P < 0.001* between groups by one-way ANOVA.

Together with IL-6, circulating IL-11 was revealed by our cytokine data as another cytokine whose level increased over time in the cxC26 group (Figure 1g). The higher expression of IL-11 was observed in Lewis lung carcinoma model, but the role of IL-11 in cancer cachexia was not studied^31^. To investigate the role of IL-11 in the C26 model, we disrupted more than 80% of the secretion of IL-11 in cxC26 cells using CRISPR/cas9 (Figure S6a). Interestingly, similar to our observations with the IL-6 KO and LIF KO, ablation of *IL-11* again slowed down the tumor growth and delayed cachexia progress (Figure S6b, c and d). Thus, IL-11 is unlikely a cachectic factor in the C26 model.

Notably, all the three cytokines IL-6, LIF, and IL-11 are members of the IL-6 family cytokines which share a four-helix bundle structure linked by loops and bind to a shared signal-transducing receptor containing the gp130 subunit^32,33^. Our results indicate their important role in the growth of the cxC26 tumor. We next focused on IL-6 and aimed to elucidate its role in tumor growth.

### IL-6 regulates tumor growth via its effects on host immune system

IL-6 family cytokines act on the JAK/STAT3 pathway for signal transduction and transcriptional regulation. In particular, the pathway is aberrantly hyperactivated in various types of tumors, and IL-6 activates the JAK/STAT3 pathway via the IL-6 receptor by an autocrine or paracrine manner, subsequently contributing to tumorigenesis^34–37^. To test whether the slowed growth of the IL-6 KO tumors results from disruption of the IL-6/JAK/STAT3 autocrine signaling pathway, we disrupted the IL-6 receptor α (*IL-6Rα*) in the cxC26 cells to block the IL-6/JAK/STAT3 autocrine signaling pathway. In contrast to the cxC26 IL-6 KO tumor, the cxC26 IL-6R*α* KOp tumor did not show any difference in tumor growth or body weight loss (Figure S7b and c). Thus, the slowed tumor growth of the cxC26 IL-6 KO tumor was independent of the IL-6/JAK/STAT3 autocrine signaling pathway.

As described earlier, the injection of one million cxC26 IL-6 KO s1 cells did not lead to any tumor growth and the injection of 10 million cxC26 IL-6 KO s1 cells showed a prolonged initiation period before starting to grow (Figure 2f). These phenomena suggest an active immune attack on the cancer cells with only a large enough initial cancer cell population escaping the attack. As it is known that tumors secrete various cytokines to regulate the tumor microenvironment^5^, we hypothesized that the cxC26 cells release IL-6 to suppress host immune response for faster growth. To test this hypothesis, we injected one million of the cxC26 cells and the cxC26 IL6 KO s1 cells into the immunodeficient NOD-SCID mice. Strikingly, the cxC26 IL-6 KO s1 tumor did not show any growth defect compared to the wild-type cxC26 tumor, in contrast to its growth on immunocompetent mice (Figure 4a and b). Thus, the cxC26-derived IL-6 promotes tumor growth by suppressing host immune response.

**Fig. 4.**
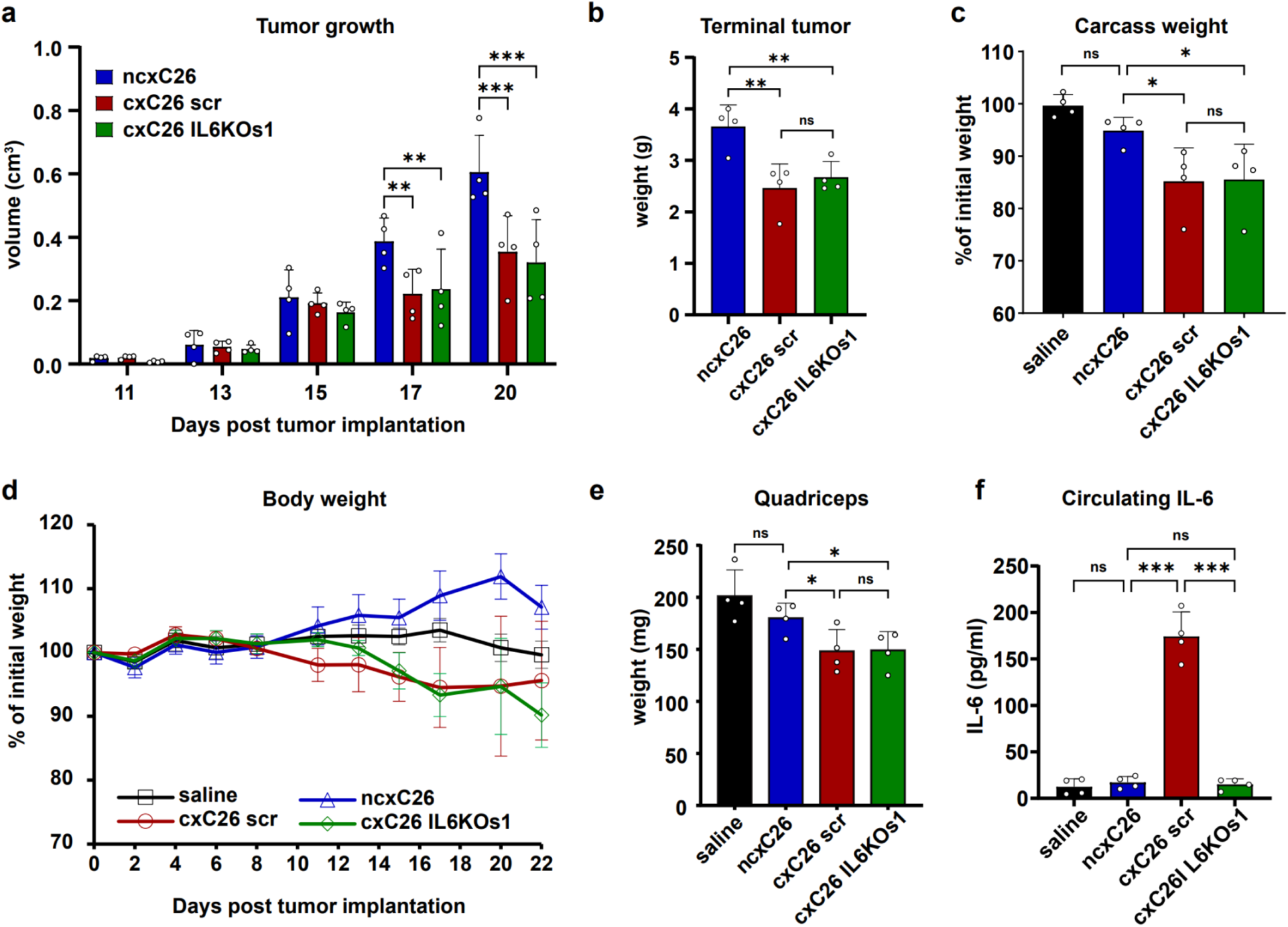
Depletion of the host immune system rescues the growth of IL-6 deficient cachectic C26 tumor. NOD-SCID male mice were inoculated with 1 × 10^6^ ncxC26 or cxC26 scr or cxC26 IL-6 KO cells. All groups were euthanized simultaneously on day 22. **a** Growth of tumors and **b** terminal tumor weight in ncxC26, cxC26 scr or cxC26 IL-6 KOs1 tumor-bearing mice. **c** The carcass weight at day 22, normalized by initial body weight. **d** Body weight **e** mass of quadriceps, and **f** concentration of circulating IL-6 at terminal time point. All data in the figure are shown as the mean ± s.d. Significance of the differences: *\*P < 0.05, ** P < 0.01, *** P < 0.001* between groups by one-way ANOVA. n = 4 per group

Notably, consistent with a previous study^23^, cachexia developed in the NOD-SCID mice bearing the cxC26 tumor (Figure 4d and e), suggesting that the lymphocytes are not mediators of cachexia. Importantly, cachexia still took place in the NOD-SCID mice bearing the cxC26 IL-6 KO s1 tumor (Figure 4d and e), without elevation of circulating IL-6 (Figure 4f), further supporting that IL-6 is not necessary for causing cachexia in this model.

### Cachectic tumor-derived IL-6 suppresses the infiltration of host immune cells

To further probe how the tumor-derived IL-6 interacts with the host immune response, we evaluated the infiltration of immune cells to the tumor. We quantified different immune cell types in tumor using flow cytometry at an early phase of tumor growth (day 8) when the cxC26 tumor and the cxC26 IL-6 KO s1 tumor had similar size and the terminal phase (day 13 for cxC26 and day 30 for cxC26 IL-6 KO s1) when both groups developed the cachexia (Figure S8). Strikingly, in the early phase of tumor growth, the proportion of leukocytes (CD45+) was 70% of all cells in the cxC26 IL-6 KO s1 tumor, 3.7-fold more abundant than that in the wild-type cxC26 tumor (Figure 5a). Furthermore, the results revealed a generally increased infiltration of individual immune cell types in the cxC26 IL-6 KO s1 tumor at the early phase of tumor growth, including B lymphocytes (CD45+/CD19+/CD11b-), macrophages (CD45+/CD11b+/F4/80+/Ly6G-), neutrophils (CD45+/CD11b+/F4/80-/Ly6G+), CD4+ T-lymphocytes (CD45+/CD3+/CD4+/CD8-) and CD8+ T-lymphocytes (CD45+/CD3+/CD4-/CD8+) (Figure 5b-f). Thus, the cxC26 tumor releases IL-6 to suppress the host immune response by inhibiting the recruitment of cancer-killing immune cells in the tumor.

**Fig. 5.**
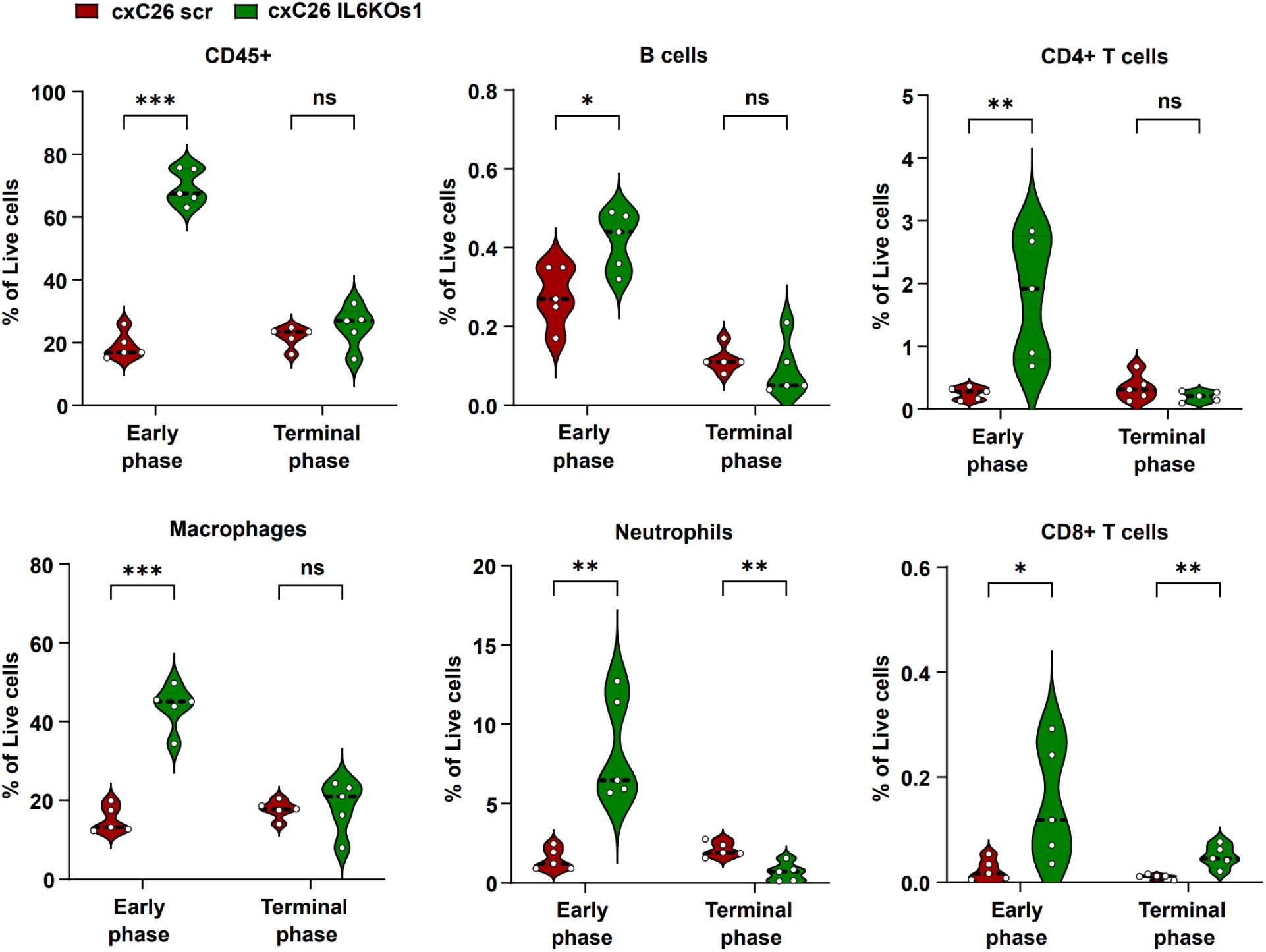
Infiltration of host immune cells into cxC26 and cxC26 IL-6 KO tumor. Mice were inoculated 1 × 10^6^ cxC26 scr or 1 × 10^7^ cxC26 IL-6 KO s1 cells. The tumor-bearing mice were euthanized at the early phase (day 8 for all groups) when they had similar tumor size and terminal phase (day 13 for cxC26 and day 30 for cxC26 IL-6 KO s1) when they developed cachexia. The tumors were dissociated into single-cell suspension and stained with surface markers. All data are presented as percent of total live cells (7-AAD negative cells) and quantitate the proportion of CD45+ cells, B cells (CD45+/CD19+/CD11b-), CD4+ T-cells (CD45+/CD3+/CD4+/CD8-), CD8+ T-cells (CD45+/CD3+/CD4-/CD8+), macrophages (CD45+/CD11b+/F4/80+/Ly6G-), and neutrophils (CD45+/CD11b+/F4/80-/Ly6G+) were enumerated using flow cytometry. PD-1 positive cells were evaluated in CD4+ and CD8+ T-cells population. Statistical significance (t-test) between groups: *0.05 < p, **0.01 < p, *** 0.001 < p. n.s., non-significance. All data in the figure are shown as the mean ± s.d. Significance of the differences: *\*P < 0.05, ** P < 0.01, *** P < 0.001* between the cxC26 scr and cxC26 IL-6 KO s1 group by Student’s t-test. n = 5 per group.

## Discussion

Identifying the cachectic factors is a major goal of cancer cachexia research. To qualify as a cachectic factor, a factor must meet the criterion that inhibition of the factor leads to less body weight loss without negatively impacting tumor progression. While body weight result is always presented as a central piece of data in studies proposing cachectic factors, it is not uncommon that tumor size and cancer status are not rigorously addressed or even ignored. In this study, we subjected one of the widely accepted cachectic factor, IL-6, to this criterion in the C26 model, and found that it did not meet the criterion as inhibiting IL-6 slowed tumor growth. We then went further and ruled out IL-6 as a cachectic factor in the C26 model by showing that cachexia still took place without elevation of circulating IL-6. Surprisingly, another widely accepted cachectic factor, LIF, was also found to affect cachexia via its effects on tumor growth in the C26 model.

In this study, we focused on the C26 model, where the role of IL-6 in cancer cachexia was originally proposed. While our results do not make any conclusions in other models of cancer cachexia where IL-6 has been implicated as a mediator of cachexia^19,38,39^, they call for closer examination of IL-6’s role in those models. In fact, in studies that reported preventive effect of cachexia by the disruption of *IL-6* gene in a pancreatic ductal adenocarcinoma (PDAC) model ^39^, a fibrosarcoma (CHX207) model^40^ and an *Apc^Min/+^* colorectal cancer model^15^, tumor growth was also reduced. More generally, our results serve as an important reminder that to establish the causality of a factor in cachexia the effect of the factor on tumor growth needs to be closely monitored.

Despite being widely cited as a cachectic factor, in cancer cachexia patients, circulating IL-6 was either not changed^41–44^, or showed modest increase (less than 3-fold)^19,45–47^, orders of magnitude lower than those measured in mouse models (up to 1,000-fold). Moreover, IL-6 is a pleiotropic cytokine, involved not only in immune and inflammatory responses but also in other processes such as exercise^48,49^ For example, in humans the circulating IL-6 level can be raised ∼1,000 times in inflammatory states^50^ and in exercise^51^. Thus, there is no strong data that supports the cachectic role of IL-6 in human cancer patients, in line with our results on IL-6 in the C26 model.

Our results revealed that IL-6 promotes the growth of the cxC26 tumor via its effect on immune response, which is in line with known roles of IL-6 in cancer proliferation^52^. Directly relevant to our results, IL-6 is known to modulate the host immune system by suppressing the adaptive immune response or recruiting tumor-associated M2 macrophages, which can promote the growth and progression of cancer^53^. In several murine cancer models, disruption of IL-6 suppressed tumor growth by restoring the anti-tumor activity of host immune system^54–56^. Specifically in the colon 26 cancer models, IL-6 level was observed to be negatively correlated with immune cell populations in tumors^16^, and mice deficient in IL-6 had increased immune cell infiltration and reduced tumor growth^57^. Thus, our observations of the increased infiltration of immune cells and slowed tumor growth in the cachectic tumor depleted of IL-6 (Figure 5) are in line with the known immunosuppression function of IL-6, though here our unique finding is that it is the tumor-derived IL-6 that represses immune response.

By invalidating the two known cachectic factors in the C26 model, our results call for the search for the real cachectic factors in the model. As demonstrated in our study, knocking out a gene in the cxC26 cells and monitoring the effects on body weight loss and tumor growth is an effective strategy to screen for cachectic genes. With the ncxC26 cell line as a control, the candidate cachectic genes are the differentially expressed genes between the cxC26 and ncxC26 tumors. The list of candidate genes can be shortened by determining whether their corresponding circulating proteins have different levels between the cxC26 and ncxC26 mouse groups. To further increase the success rate of this strategy, we propose the use of the cxC26 IL-6 KO cell line together with the NOD SCID mice as a “cleaner” model. First, the cxC26 IL-6 KO cell line does not produce IL-6, which is a pleiotropic cytokine that exerts multiple functions including potentially regulating gene expression in the tumor. Second, the immunodeficient mice are free of adaptive immune responses, which can result in altered gene expression in the tumor. Thus, though in principle the cachectic factors can be any molecules such as metabolites or lipids, this approach will potentially reveal tumor-derived proteins that contribute to cachexia.

## Material and Method

### Animals

Animal care and experimental procedures were conducted with the approval of the Institutional Animal Care and Use Committees (IACUC) of Harvard Medical School and Harvard T.H. Chan School of Public Health. CD2F1, Balb/c and NOD-SCID male mice were purchased from Charles River and used for this study when the mice were 13 – 18 weeks old. C26 cells were harvested and implanted in the right flank of mice described in a previous paper^28^. All mice were housed (3-4 per cage) under regular light-dark cycles of 12 hours with ad libitum of standard chow diet (PicoLab 5053, LabDiet) and water.

### Cell culture

The cachectic C26 (cxC26) and non-cachectic C26 (ncxC26) cells were a kind gift from Andrea Bonetto^28^ and Nicole Beauchemin, respectively. Nicole Beauchemin obtained the ncxC26 from Brattain MG, who established C26 (also named CT26 or colon carcinoma 26) cell line^58^. The C26nci (C26 from National Cancer Institute) and C26nci LIF KO were a kind gift from Robert Jackman^29^. All C26 cells were maintained using high glucose DMEM (Corning) with 10% fetal bovine serum (FBS, R&D Systems) and 1% penicillin/streptomycin (P/S, Hyclone).

### Construction of knock-out cells

To construct the knock-out (KO) cells, two different sets of single guide RNA (sgRNA) targeting the indicated gene was cloned into the BsmBI site of the lentiCRISPRv2 vector (Addgene #52961) as previous described^59^.The sequences of the sgRNA for each target genes are shown in Supplementary Table 1. For lentivirus production, HEK293T cells were plated in DMEM (10% FBS) and transfected two sets of sgRNA-containing lentiviral vector plasmids along with pMD2.G (Addgene #12259) and psPAX2 (Addgene #12260) into the HEK293T cells using Lipofectamine 3000 (Thermo Fisher Scientific). After 24 hours of incubation, the lentivirus-containing medium was collected and filtered through a 0.45 µm syringe filter. The cxC26 cells were then cultured with the lentivirus-containing medium in the presence of polybrene (5 µg/ml, Sigma-Aldrich) for 24 hours. Transfected cells were subsequently selected in the presence of puromycin (3 ug/ml, Santa Cruz Biotechnology) in fresh DMEM (+10% FBS and 1% P/S) media. To obtain clonal cell lines of cxC26 IL-6 KO cells, the cxC26 IL-6 KO pool cells were plated onto 96 wells by serial dilution. Two single cells were isolated and expanded in medium containing puromycin. Two subclones were named as cxC26 IL-6 KO subclone 1 (s1) and cxC26 IL-6 KO subclone 2 (s2), respectively.

### Analysis of cancer cachexia phenotype

To obtain the phenotype of cancer cachexia, the body weight and tumor size were monitored every other day post-tumor implantation. For tumor volume calculation, the length and width of the tumor were measured using a caliper. The tumor volume was obtained using the formula V = (W(2) × L)/2. The mice were euthanized when they developed more than 15% of body weight loss compared to the initial body weight. To assess body composition, lean mass and fat mass of live mice were measured using an EchoMRI-100 (EchoMRI LLC). To measure the grip strength, mice were allowed to grab the metal grid and were then pulled backwards by an experimenter until the grasp was released. The peak tension was measured using a Grip Strength meter (Bioseb).

### Cytokines measurement

Circulating cytokines were measured in mouse plasma using the Luminex 44-cytokines panel by Eve Technologies (Calgary, Canada). Circulating LIF and IL-11 were also measured using Mouse LIF Quantikine ELISA Kit (R&D Systems) and IL-11 SimpleStep ELISA kit (abcam), respectively. And circulating IL-6 was also determined using high sensitive IL-6 Mouse ProQuantum qPCR Immunoassay kit (Invitrogen) with real-time PCR.

### Flow cytometry

CD2F1 male mice were inoculated with 1 × 10^6^ cxC26 cells or 1 × 10^7^ cxC26 IL-6 KO s1 cells, and the tumors were harvested at day 8 or at the terminal point when tumor-bearing mice developed cachexia (n = 5). To prepare single-cell suspension, the harvested tumor was immediately minced using a razor blade in 1 ml of 10% FBS containing RPMI (Gibco), added 2 ml of ice-cold HBSS (Fisher Scientific) containing collagenase I (10 U/ml, Worthington Biochemical), collagenase IV (400 U/ml, Worthington Biochemical), and DNase I (30 U/ml, STEMCELL Technology), and incubated with rotation for 25 min at 37℃ in hybridization oven. Finally, the digests were gently crushed with a pestle and filtered through a 70-μm cell strainer (BD Falcon) to yield single-cell suspensions. To remove red blood cells, the suspension was treated with RBC lysis buffer (BioLegend) on ice for 2 min. The cells were then washed with RPMI media and resuspended in Cell staining buffer (BioLegend). For each sample, 1 × 10^6^ cells were treated with TruStain FcX^tm^ Plus (anti-mouse CD16/32, Biolegend) for 10 min and labeled with various fluorescent antibodies. The antibodies for flow cytometry are listed in Supplementary Table 2. The dead cells were excluded by 7-AAD staining (BioLegend). The labeled single cells were analyzed using flow cytometry (BD LSR Fortessa), and Flowjo software (Tree Star Inc.) was used for data analysis.

## Statistical analysis

All statistical analyses for murine data were performed in GraphPad Prism 9 or R studio. Quantitative data are reported as mean with standard deviation (s.d.). Two-group comparisons were analyzed by a two-tailed Student’s t-test and more than two-group comparisons were analyzed by One-way ANOVA or Two-way ANOVA. Adjusted p-values obtained for multiple comparisons in R using the Benjamini–Hochberg method. For all analyses, a p-value of < 0.05 was considered significant (*p < 0.05, **p < 0.01, and ***p < 0.001). The R package ComplexHeatmap was used to generate the heatmap for the concentration of cytokines.

## Data availability

All data generated or analyzed during this study are included in this published article (and its supplementary figures and tables). Source data are provided with this paper.

## Acknowledgements

We would like to thank Dr. Andrea Bonetto (cxC26), Dr. Nicole Beauchemin (ncxC26), and Dr. Robert Jackman (C26nci and C26nci LIF KO) for graciously providing cell lines used for our studies. We are grateful to Dr. Tobias Janowitz, Dr. Eileen White, Dr. Marcus Goncalves, and Dr. Jessalyn Ubellacker for valuable suggestions and critical reading of the manuscript. We thank Nhien Tran for technical support with the construction of the cxC26 IL-6Ra KO cells. This work was delivered as part of the CANCAN team supported by the Cancer Grand Challenges partnership funded by Cancer Research UK (CGCATF-2021/100022) and the National Cancer Institute (1 OT2 CA278685-01). This work was also supported by R00DK117066 (S.H.).

## Author contributions

Y.Y.K. and S.H. designed experiments and wrote the manuscript. Y.Y.K., performed experiments and data analysis.

## Supplementary Figures

**Supplementary Fig. 1.**
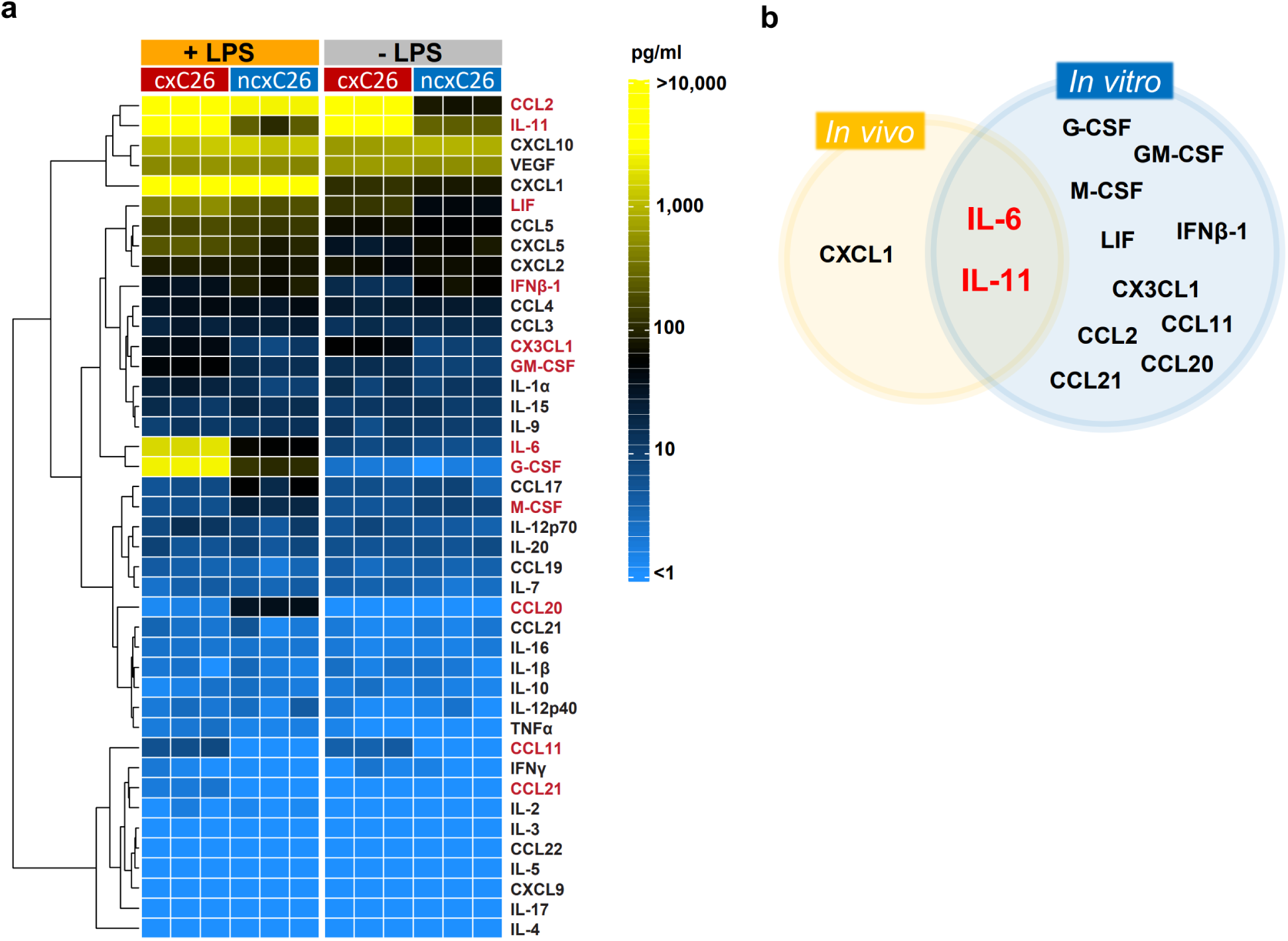
Profiling of secretory cytokines from cxC26 and ncxC26 cells. **a** Levels of cytokines in conditioned media with or without LPS treatment for 24 hours. The measurement was done using the Luminex 44 cytokines panel. Cytokines highlighted in red are those with significantly altered levels under LPS treatment between cxC26 and ncxC26 (adjusted p-value < 0.01 by Student’s t-test, n = 3 per group). **b** Comparison between significantly changed cytokines by cxC26 in *in vivo* (Figure 1g) and *in vitro*.

**Supplementary Fig. 2.**
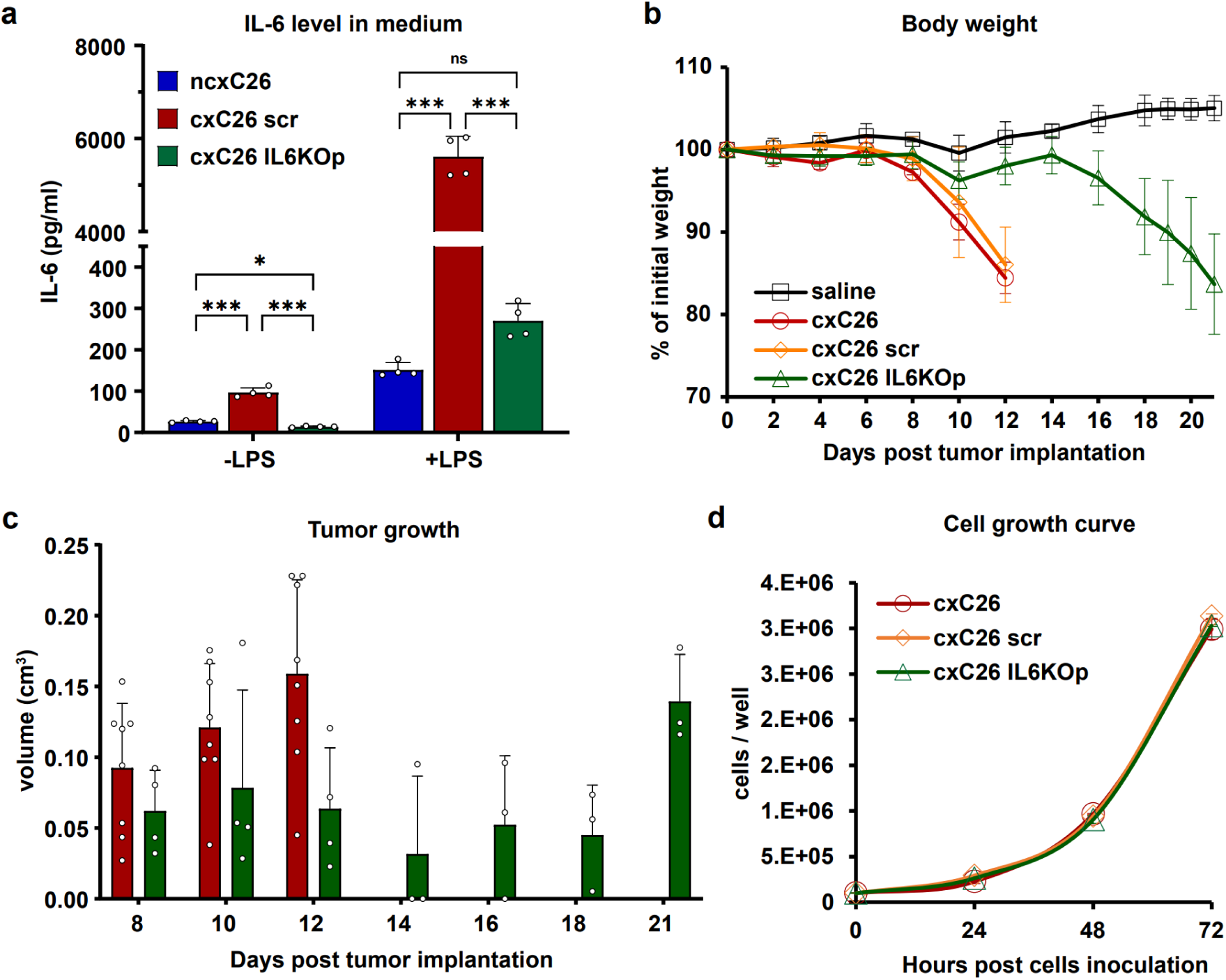
Characterization of cxC26 IL-6 KO pool (IL6KOp) **a** Concentration of IL-6 in conditioned media of cxC26 scr (CRISPR/Cas9 scrambled gRNA control) and cxC26 IL-6 KOp with or without treatment of lipopolysaccharide (LPS) for 24 hours. **b** Body weight and **c** tumor volume post-tumor implantation. CD2F1 male mice were inoculated with 1 × 10^6^ cxC26 or cxC26 IL-6 KOp cells. n = 4 per group. **d** Growth curve of cxC26, cxC26 scr and cxC26 IL-6 KOp cells in *in vitro.* n = 3 per group. Significance of the differences: *\*P < 0.05, ** P < 0.01, *** P < 0.001* between groups by one-way ANOVA.

**Supplementary Fig. 3.**
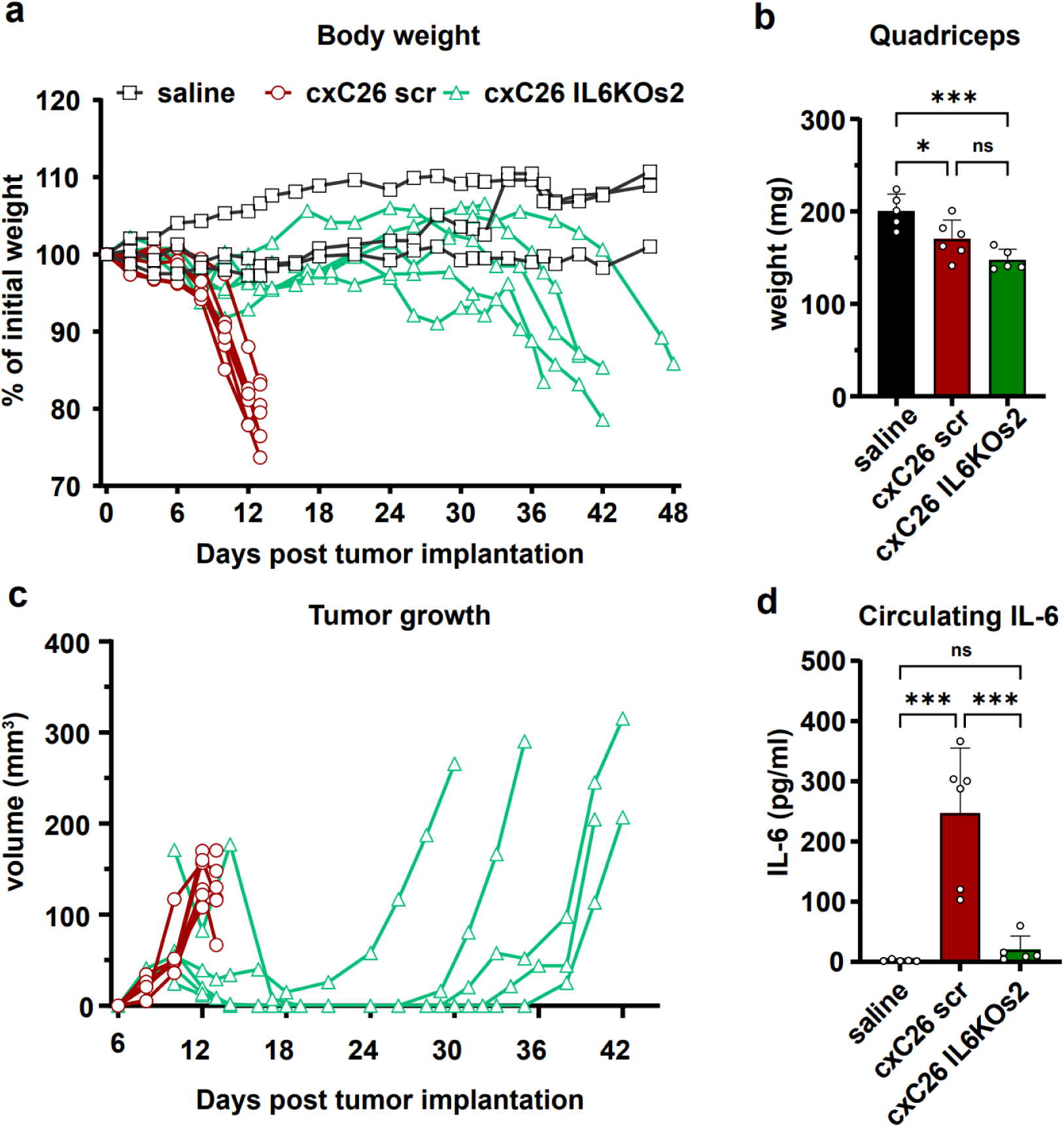
Characterization of cxC26 IL-6 KO subclone 2 (s2) CD2F1 male mice were inoculated with 1 × 10^6^ cxC26 scr or 1 × 10^7^ cxC26 IL-6 KO s2 cells. **a** Body weight, **b** terminal muscle mass, and **c** circulating IL-6 concentration at the end of experiment when the tumor-bearing animals developed cachexia (more than 15% of body weight loss). Significance of the differences: *\*P < 0.05, ** P < 0.01, *** P < 0.001* between groups by one-way ANOVA. n = 5 for saline and cxC26 IL-6 KO s2, n = 6 for cxC26.

**Supplementary Fig. 4.**
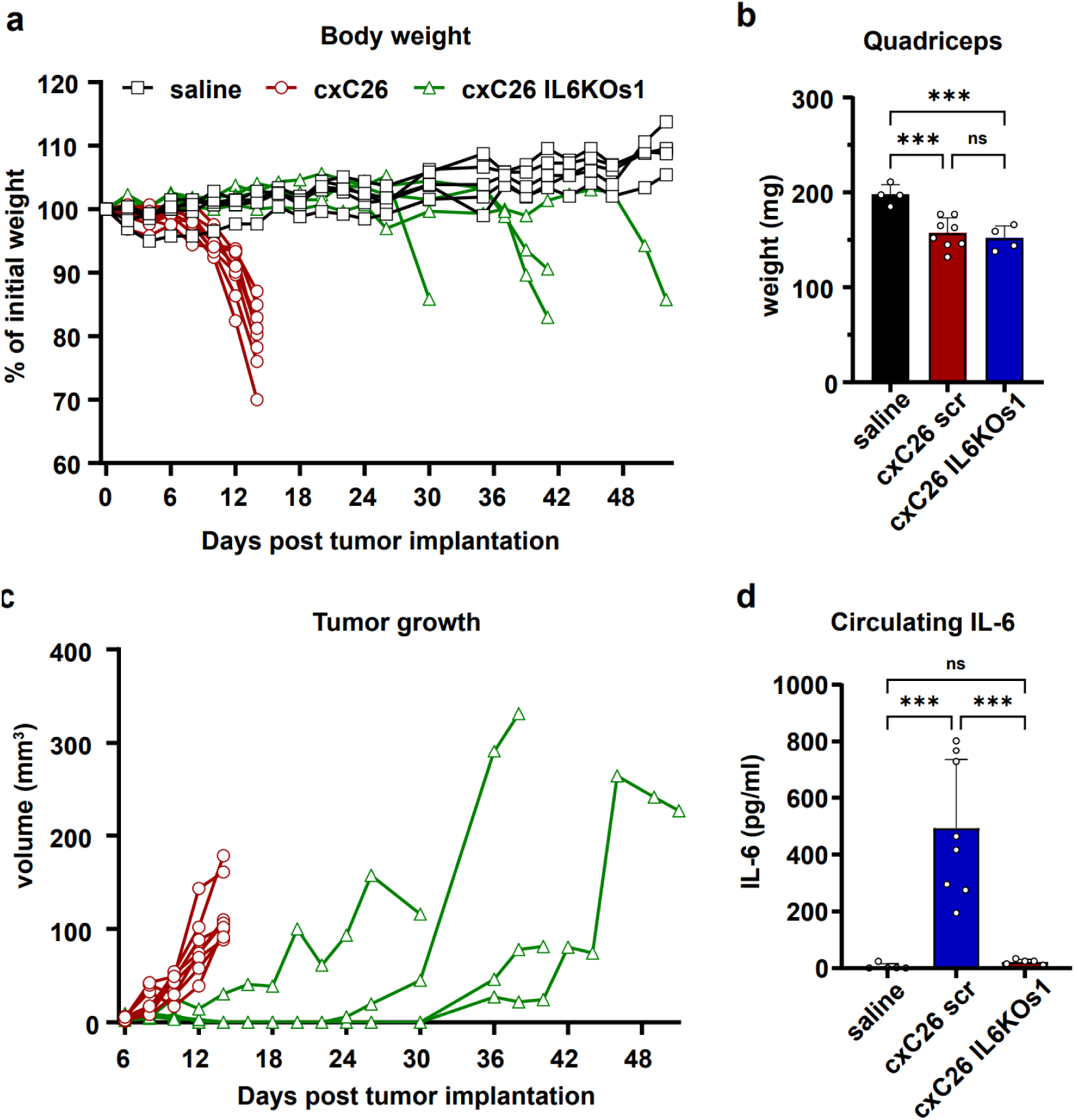
Validation of body weight loss and slow growth of C26 IL-6 KO subclone 1 (s1) tumor in Balb/c mice. Balb/c male mice were inoculated with 1 × 10^6^ cxC26 scr or 1 × 10^7^ cxC26 IL-6 KO s1 cells. **a** Body weight, **b** terminal muscle mass and **c** circulating IL-6 concentration at the end of experiment when the tumor-bearing animals developed cachexia (more than 15% of body weight loss). Significance of the differences: *\*P < 0.05, ** P < 0.01, *** P < 0.001* between groups by one-way ANOVA.

**Supplementary Fig. 5.**
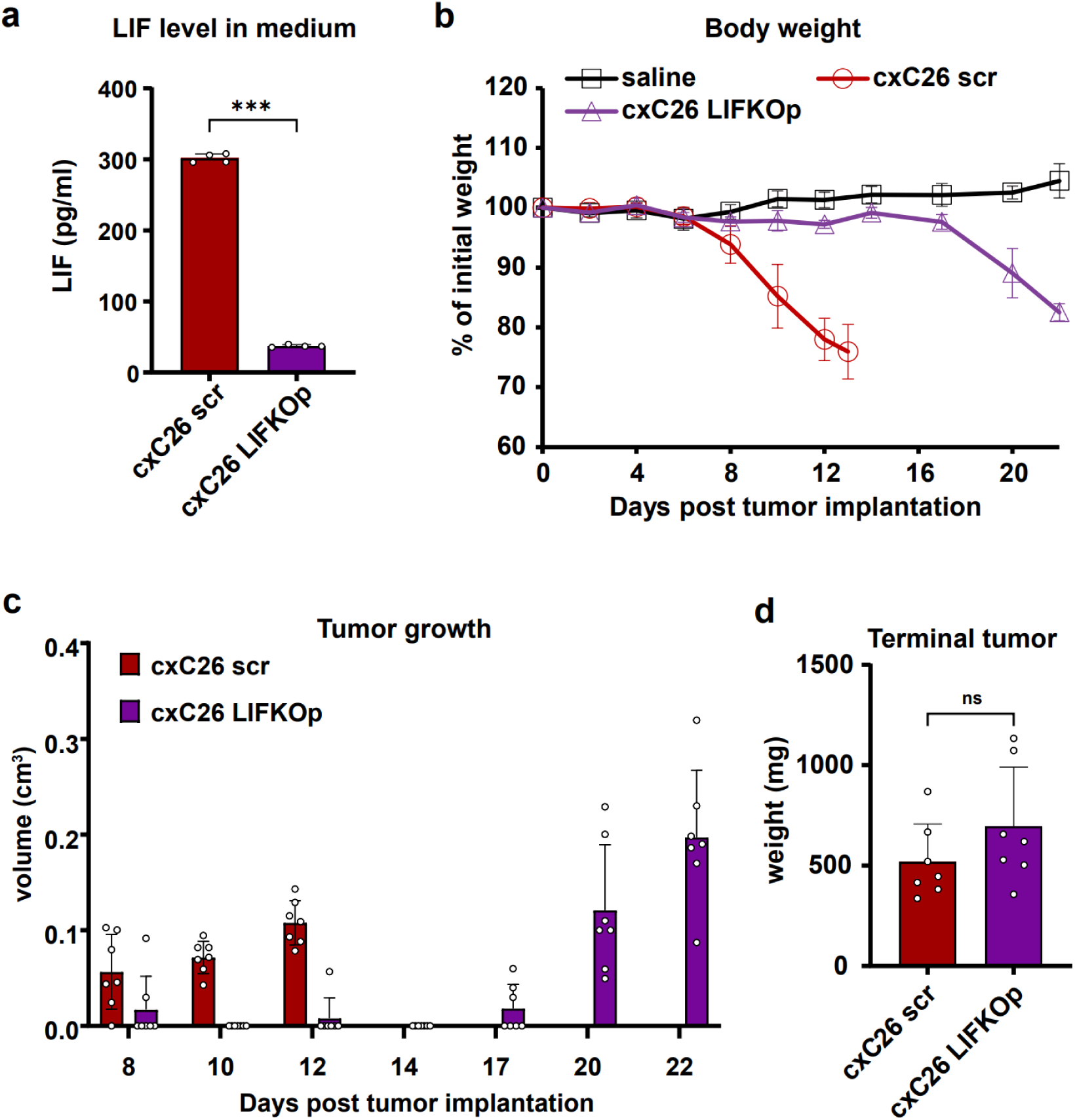
Effects of *LIF* knockout in the cxC26 tumor. **a** Concentration of LIF in conditioned media of cxC26 scr (CRISPR/Cas9 scrambled gRNA control) and cxC26 LIF KOp. **b** Body weight, **c** tumor growth and **d** tumor mass at the end of experiment when the tumor-bearing animals developed more than 15% of body weight loss compared to their initial body weight. CD2F1 male mice were inoculated with 1 × 10^6^ cxC26 scr or xC26 LIF KOp cells. Significance of the differences: *\*P < 0.05, ** P < 0.01, *** P < 0.001* between groups by Student’s t-test. n = 5 for saline, n = 7 for cxC26 scr, n = 7 for cxC26 LIF KOp.

**Supplementary Fig. 6.**
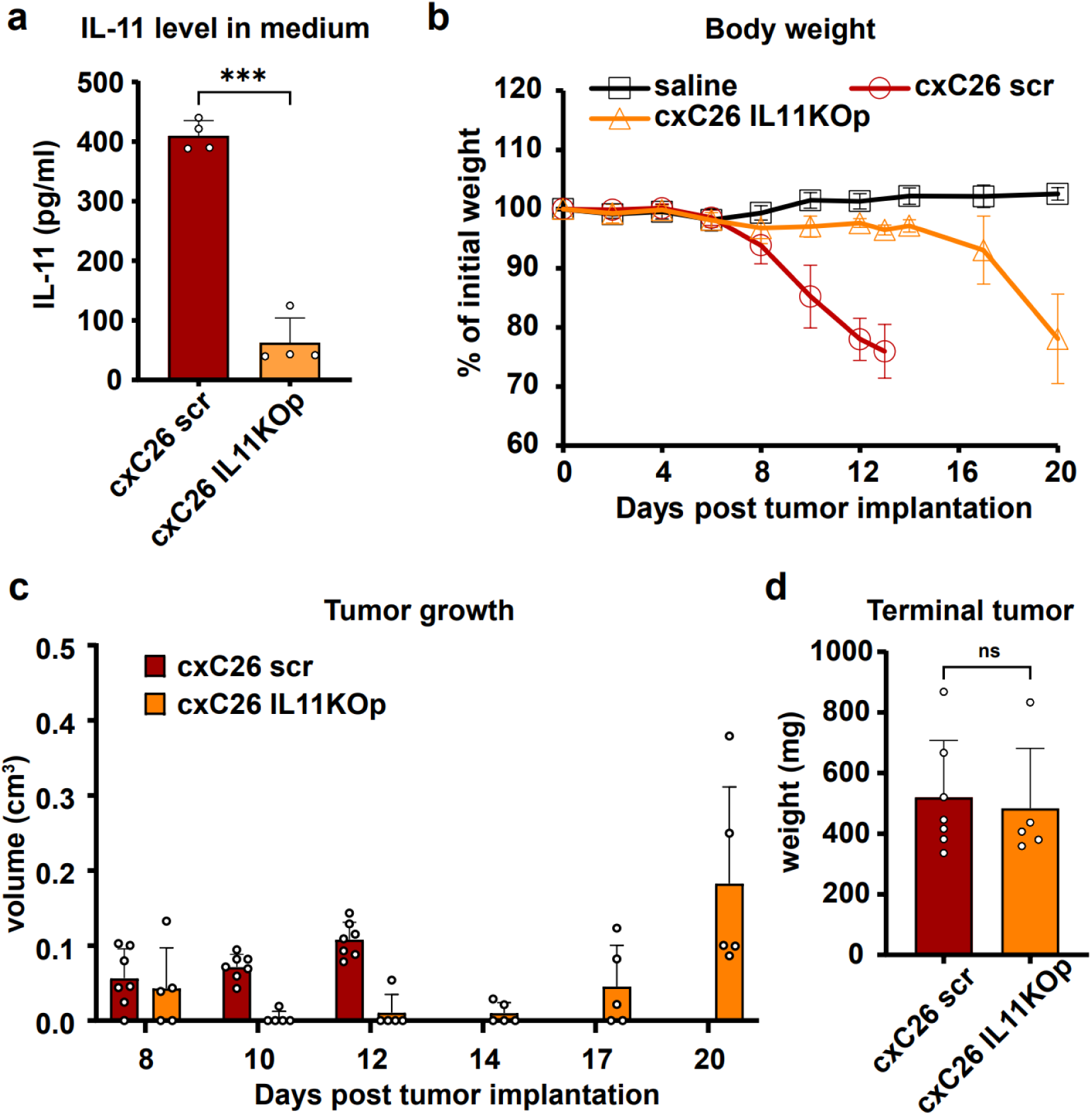
Characterization of *IL-11* knockout in the cxC26 tumor. **a** Concentration of LIF in conditioned media of cxC26 scr (CRISPR/Cas9 scrambled gRNA control) and cxC26 IL-11 knockout pool (KOp). **b** Body weight, **c** tumor growth, and **d** tumor mass at the end of experiment when the tumor-bearing animals developed more than 15% of body weight loss compared to their initial body weight. CD2F1 male mice were inoculated with 1 × 10^6^ cxC26 scr or xC26 IL-11 KOp cells. Significance of the differences: *\*P < 0.05, ** P < 0.01, *** P < 0.001* between groups by Student’s t-test. n = 5 for saline, n = 7 for cxC26 scr, n = 5 for cxC26 IL-11 KOp.

**Supplementary Fig. 7.**
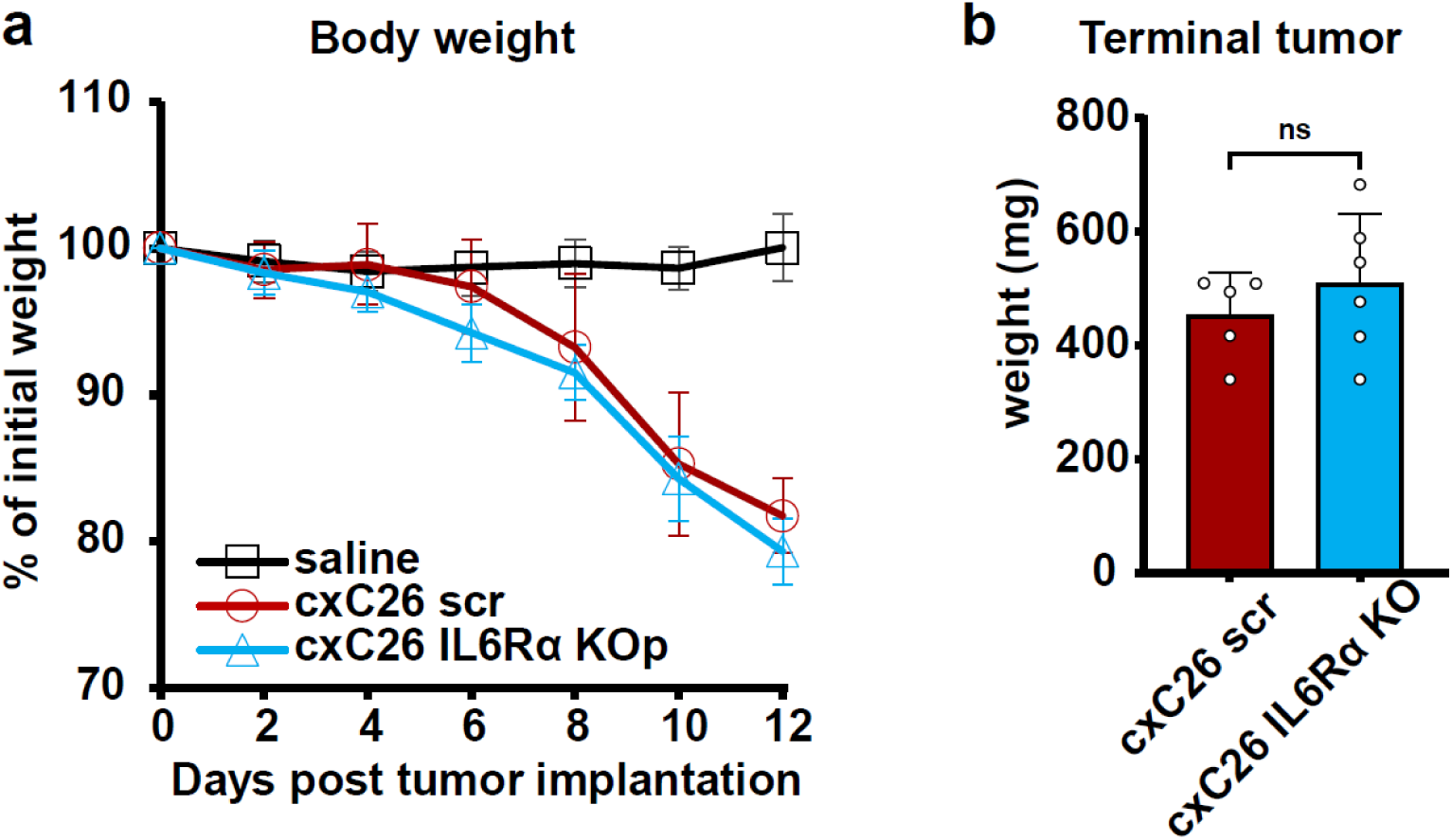
Inhibition of autocrine IL-6 signaling pathway by disrupting IL-6 receptor in cxC26 cells. cxC26 IL-6 receptor α (*IL-6Rα*) knock-out pool (KOp) was constructed using CRISPR/cas9. CD2F1 male mice were inoculated with 1 × 10^6^ cxC26 or cxC26 IL-6Rα KOp cell. **a** Body weight curve and **b** terminal tumor weight at day 12. Significance of the differences: *\*P < 0.05, ** P < 0.01, *** P < 0.001* between groups by Student’s t-test. n = 4 for saline, n = 5 for cxC26 scr, n = 6 for cxC26 IL-6Rα KOp.

**Supplementary Fig. 8.**
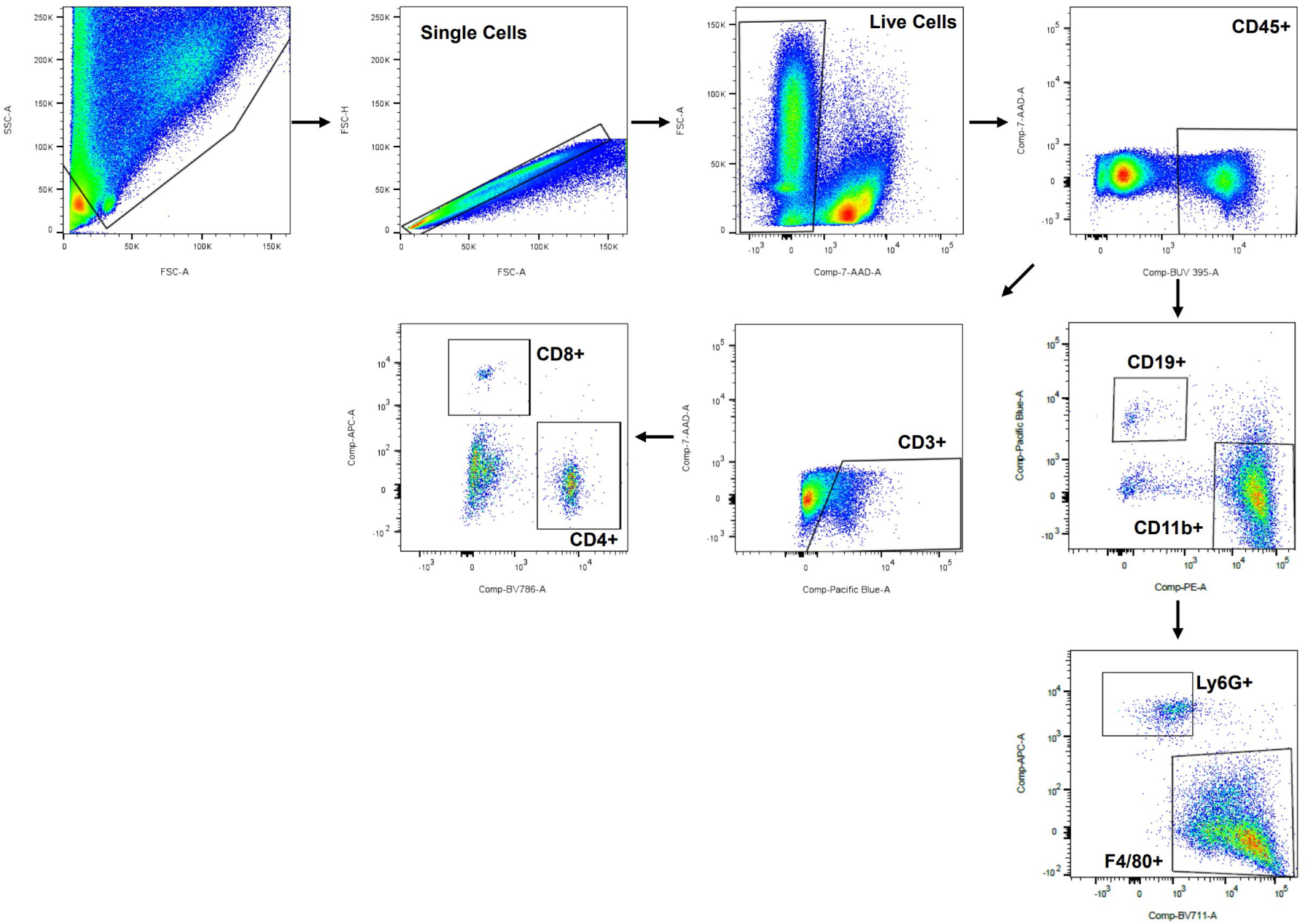
Representative gating strategy for the evaluation of immune cell populations in cxC26 scr and cxC26 IL-6 KO s1 tumors. Tumors were harvested and dissociated into single-cell suspension and stained with various surface markers which are described in Supplementary Table 2.

## Supplementary Tables

**Supplementary Table 1.**
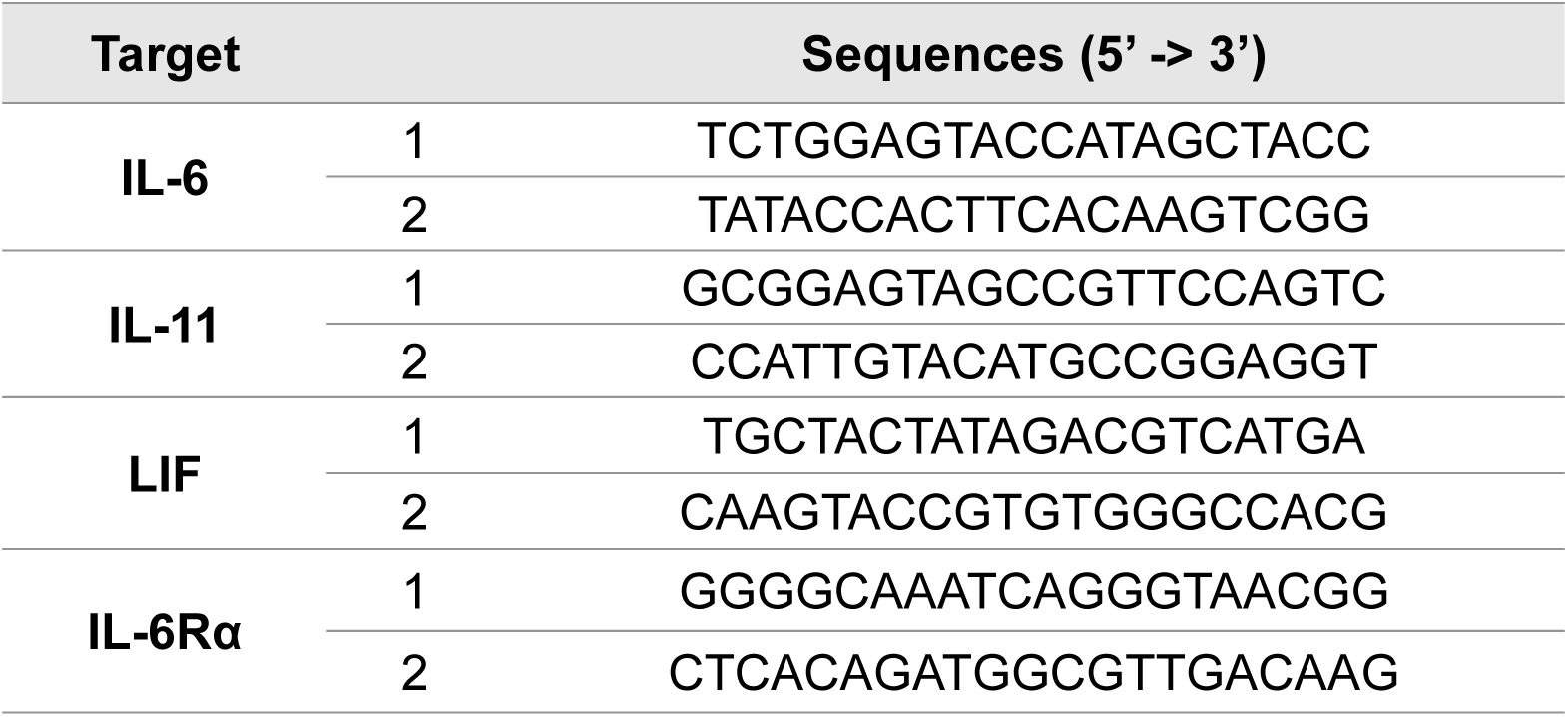
The list of guide RNA sequences designed for CRISPR/Cas9 constructs.

**Supplementary Table 2.**
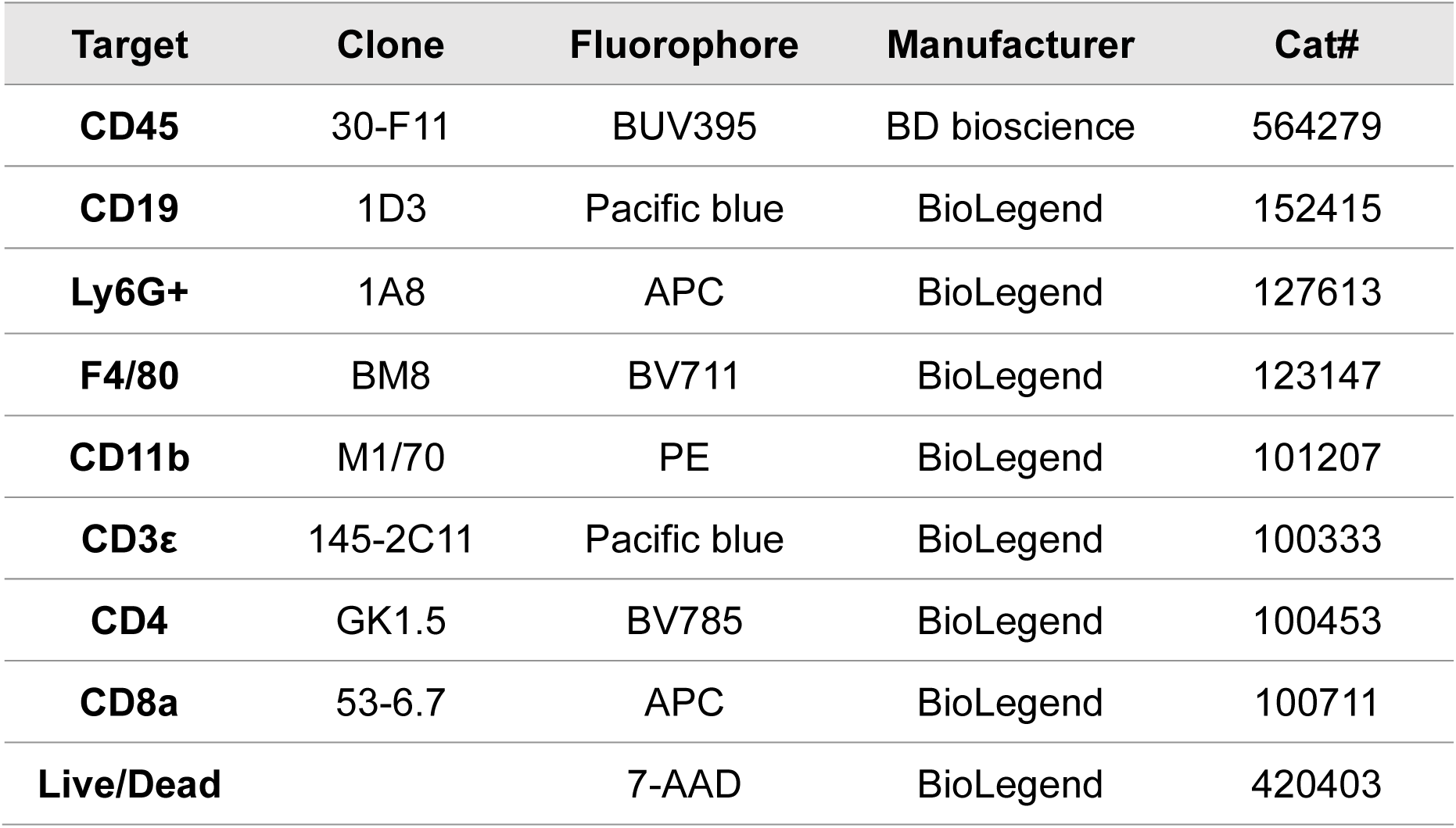
Materials used for flow cytometry.

| Target | Clone | Fluorophore | Manufacturer | Cat# |
| --- | --- | --- | --- | --- |
| CD45 | 30-F11 | BUV395 | BD bioscience | 564279 |
| CD19 | 1D3 | Pacific blue | BioLegend | 152415 |
| Ly6G+ | 1A8 | APC | BioLegend | 127613 |
| F4/80 | BM8 | BV711 | BioLegend | 123147 |
| CD11b | M1/70 | PE | BioLegend | 101207 |
| CD3ε | 145-2C11 | Pacific blue | BioLegend | 100333 |
| CD4 | GK1.5 | BV785 | BioLegend | 100453 |
| CD8a | 53-6.7 | APC | BioLegend | 100711 |
| Live/Dead |  | 7-AAD | BioLegend | 420403 |

